# Alternative polyadenylation drives oncogenic gene expression in pancreatic ductal adenocarcinoma

**DOI:** 10.1101/752295

**Authors:** Swati Venkat, Arwen A. Tisdale, Johann R. Schwarz, Abdulrahman A. Alahmari, H. Carlo Maurer, Kenneth P. Olive, Kevin H. Eng, Michael E. Feigin

## Abstract

Alternative polyadenylation (APA) is a gene regulatory process that dictates mRNA 3’-UTR length, resulting in changes in mRNA stability and localization. APA is frequently disrupted in cancer and promotes tumorigenesis through altered expression of oncogenes and tumor suppressors. Pan-cancer analyses have revealed common APA events across the tumor landscape; however, little is known about tumor type-specific alterations that may uncover novel events and vulnerabilities. Here we integrate RNA-sequencing data from the Genotype-Tissue Expression (GTEx) project and The Cancer Genome Atlas (TCGA) to comprehensively analyze APA events in 148 pancreatic ductal adenocarcinomas (PDAs). We report widespread, recurrent and functionally relevant 3’-UTR alterations associated with gene expression changes of known and newly identified PDA growth-promoting genes and experimentally validate the effects of these APA events on expression. We find enrichment for APA events in genes associated with known PDA pathways, loss of tumor-suppressive miRNA binding sites, and increased heterogeneity in 3’-UTR forms of metabolic genes. Survival analyses reveal a subset of 3’-UTR alterations that independently characterize a poor prognostic cohort among PDA patients. Finally, we identify and validate the casein kinase CK1α as an APA-regulated therapeutic target in PDA. Knockdown or pharmacological inhibition of CK1α attenuates PDA cell proliferation and clonogenic growth. Our single-cancer analysis reveals APA as an underappreciated driver of pro-tumorigenic gene expression in PDA via the loss of miRNA regulation.

## INTRODUCTION

Pancreatic ductal adenocarcinoma (PDA) is a lethal cancer with a 5-year survival rate of 9%^1^. Extensive sequencing studies have uncovered recurrently mutated genes (*KRAS, TP53, SMAD4, CDKN2A*) and dysregulated pathways (axon guidance, cell adhesion, small GTPase signaling, protein metabolism) driving disease initiation and progression^2–4^. Gene expression profiles from hundreds of patient samples have allowed the identification of several PDA subtypes, with implications for treatment response and patient outcome^5–10^. Gene expression can be dysregulated in cancer through a variety of mechanisms, including genomic amplification/deletion, epigenetic modification and noncoding mutations in promoters/enhancers^11–15^. For example, recurrent noncoding mutations in PDA are enriched in promoters of cancer-associated genes and pathways^16^. However, our understanding of the mechanisms driving dysregulated gene expression in cancer remains incomplete. Determining the regulatory mechanisms driving dysregulated gene expression is critical to understanding disease pathogenesis. One such regulatory mechanism that has recently gained recognition as a critical driver of gene expression is alternative polyadenylation (APA).

APA is a post-transcriptional process that generates distinct mRNA isoforms of the same gene as a mechanism to modulate gene expression. This includes transcripts that have identical coding sequences but vary only in their 3’-UTR length^17–19^. Changes in 3’-UTR length can modulate mRNA stability, function or subcellular localization through disruption of miRNA or RNA-binding protein regulation^18,20,21^. APA is driven by a large complex of polyadenylation factors that recognize a series of highly conserved sequences within the 3’-UTR on the newly synthesized pre-mRNA before cleavage and addition of the poly(A) tail^18,22,23^. As most transcripts contain multiple polyadenylation sites (PAS), the choice of where to cleave is a critical determinant of 3’-UTR length. In humans, a majority of genes (51-79%) express alternative 3’-UTRs, demonstrating the widespread nature of this process^24^. Indeed, APA has important roles in muscle stem cell function, cell proliferation, chromatin signaling, pluripotent cell fate, cellular senescence and other physiological processes^25–29^. Recently, dysregulation of APA has gained recognition as a driver of tumorigenesis^28,30–33^. APA factor expression is altered in a variety of cancer types and promotes tumorigenesis by regulating the expression of oncogenes (via loss of miRNA regulation) and tumor suppressors (via disruption of competing-endogenous RNA crosstalk)^32–36^. The relevance of APA in cancer was established with the discovery of a systemic increase in the usage of a proximal PAS leading to consistently shortened 3’-UTRs of oncogenes such as Insulin-like growth factor 2 mRNA-binding protein 1 (*IMP1*), Ras-Related C3 Botulinum Toxin Substrate 1 (*RAC1*) and *Cyclin D2*^30,33^. Functional studies of the genes comprising the APA machinery have highlighted their relevance to tumor growth; for example, in glioblastoma, overexpression of the APA factor *NUDT21* (a repressor of proximal 3’-UTR PAS usage) reduces tumor cell proliferation and inhibits tumor growth *in vivo*^32^. Subsequently, a number of pan-cancer analyses have utilized standard RNA-sequencing (RNA-seq) data to identify 3’-UTR shortening and lengthening events across cancer types^37–41^. While these analyses have uncovered recurrent APA events across multiple tumor types, they also detected tumor type-specific events^42^. Additionally, differential 3’-UTR processing has been shown to drive tissue-specific gene expression^43^. However, there has been no in-depth single cancer analysis with a sufficiently large patient cohort to unravel disease-specific APA alterations. Furthermore, none of the pan-cancer studies have included PDA due to a lack of matched normal controls and therefore, the landscape of APA in PDA remains completely uncharacterized.

To determine the relevance of APA in PDA, we performed a comprehensive analysis of the changes in PAS usage using RNA-seq data from 148 PDA tumors from The Cancer Genome Atlas TCGA-PAAD (Pancreatic Adenocarcinoma) study and 184 normal pancreata from the Genotype-Tissue Expression (GTEx) project^44,45^. We performed a systems level analysis to identify trends in APA, impacts on gene expression, and effects of miRNA regulation. We discovered widespread 3’-UTR shortening events in PDA, including a subset of 68 genes shortened in >90% of patients. These 3’-UTR shortened genes did not overlap with commonly mutated PDA genes, but were enriched in PDA pathways. Furthermore, we found preferential loss of known tumor suppressive miRNA binding sites within the shortened 3’-UTRs, suggesting that APA may be acted upon by selection during tumor progression. Importantly, we identify a subset of 20 genes that detect a poor outcome cohort in PDA patients, highlighting the prognostic power of APA. Experimental validation revealed APA as a novel mechanism of regulation for known PDA growth-promoting genes. Furthermore, using computational, pharmacological and genetic approaches, we identified the casein kinase CK1α as a new therapeutic target in PDA. Our in-depth analysis reveals APA as a recurrent, widespread mechanism underlying oncogenic gene expression changes through loss of tumor suppressive miRNA regulation in pancreatic cancer.

## RESULTS

To analyze differences in APA profiles between tumor and normal samples, we selected 148 patients out of the total 178 PDA patients with aligned RNA-seq data from the TCGA-PAAD study. We excluded 30 patients in the cohort that did not have histologically observable PDA tumors^4^. Due to the paucity of RNA-seq data from matched normal tissues within the TCGA-PAAD study, we procured raw RNA-seq reads from 184 normal pancreata from the GTEx project. The library preparation and sequencing platform were identical for the TCGA-PAAD study and GTEx pancreata data^4,45^, thereby minimizing potential batch effects. Several previous studies have successfully compared TCGA and GTEx gene expression data, noting minimal batch effects when processed in an identical manner^46–48^. Therefore, these datasets were processed identically and analyzed for differences in APA in our downstream analyses (Supp. Fig. 1). To allow a rigorous comparison between GTEx normal pancreas and TCGA-PAAD tumor samples, we aligned raw reads from the GTEx RNA-seq data as per the TCGA pipeline. We processed the tumor and normal aligned files to generate coverage files that were used as an input for the DaPars (Dynamic Analysis of Alternative Polyadenylation from RNA-Seq) algorithm^41^. DaPars is a regression-based algorithm that performs de-novo identification of APA events between two conditions using standard RNA-seq data^32,33,41^. DaPars generates a mean PDUI score (Percentage Distal Usage Index) for each gene, quantifying the extent of usage of the distal PAS across each group. Genes favoring distal PAS usage (long 3’-UTRs) have PDUI scores near 1, while genes favoring proximal PAS usage (short 3’-UTRs) have PDUI scores near 0. A change in the mean PDUI score between tumor and normal samples for each gene (ΔPDUI) is then calculated and used to indicate tumor-associated 3’-UTR shortening (ΔPDUI < 0) or lengthening (ΔPDUI > 0) events.

### Integrative analysis of GTEx and TCGA-PAAD RNA-seq data identifies 3’-UTR shortening events associated with PDA

To determine the extent of APA-mediated 3’-UTR shortening and lengthening in PDA, we compared the PDUI scores for each gene between the tumor and normal samples (Fig. 1A,B). While the majority of genes do not undergo changes in APA, PDA patients are characterized by a greater number of significant 3’-UTR shortening events (red dots, n=271) as compared to significant lengthening events (blue dots, n=191) (Fig. 1B). A higher number of 3’-UTR shortening events compared to lengthening events in PDA is consistent with patterns observed in multiple pan-cancer analyses^30,41,49^. The tumor-associated shortening and lengthening events were predominantly 100-300bp and 200-300bp in length, respectively (Fig. 1C). Amongst the genes found to have significantly shortened 3’-UTR lengths were many known PDA growth-promoting genes, including *PAF1* (Polymerase Associated Factor 1), *FLNA* (Filamin-A), *ENO1* (α Enolase), *RALGDS* (Ral guanine nucleotide dissociation stimulator), *TRIP10* (Thyroid Hormone Receptor Interactor 10) and *ALDOA* (Aldolase A). *ALDOA* and *PAF1* have recently been described as oncogenes in PDA^50–53^, while *ENO1, RALGDS, TRIP10* and *FLNA* are known to mediate pancreatic cancer cell proliferation, survival and migration^54–59^. We did not detect 3’-UTR alterations in recurrently mutated PDA genes, reflecting the predominant role of APA in regulating gene expression rather than gene function. We visualized the 3’-UTR profiles of these genes between TCGA and GTEx samples to confirm 3’-UTR shortening (see *FLNA*, *PAF1* as examples, Fig. 1D).

**Figure 1.**
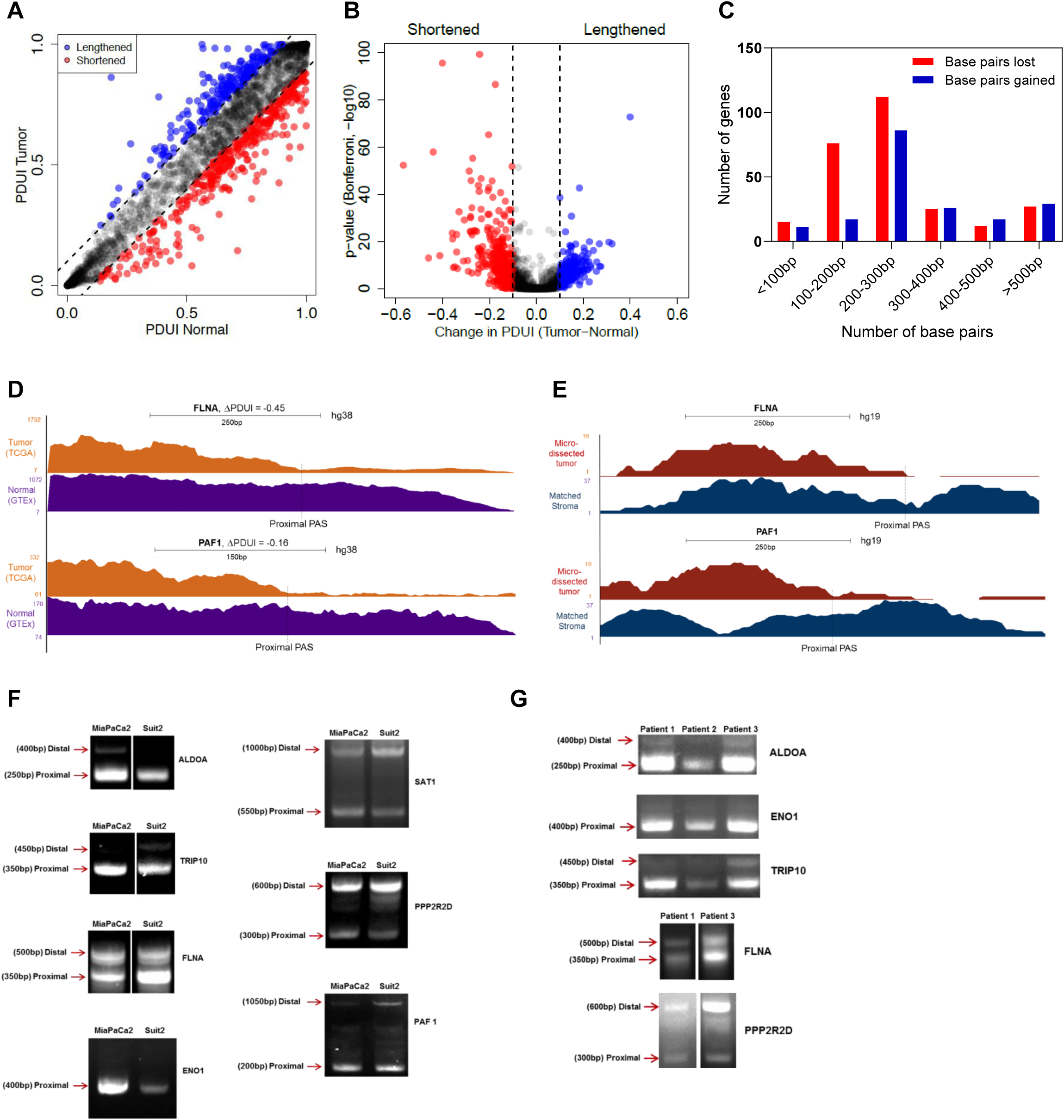
Integrative analysis of RNA-seq data identifies 3’-UTR alterations associated with PDA. (A) A plot of PDUI score of each gene in human tumor and normal samples. Dashed lines represent 0.1 cutoffs. Blue dots represent 3’-UTR lengthened genes while red dots represent 3’-UTR shortened genes. (B) A volcano plot denoting 3’-UTR shortened (red) and lengthened (blue) gene hits (FDR<0.01) whose |ΔPDUI| > 0.1. (C) A plot showing the number of base pairs lost/gained by 3’-UTR altered genes. (D) UCSC genome browser plot depicting the 3’-UTR RNA-seq density profile of two 3’-UTR shortened genes (*FLNA* and *PAF1*) to highlight the coverage differences between tumor (orange) and normal (purple) patient samples. (E) UCSC genome browser plot highlighting the 3’-UTR profile differences between *FLNA* and *PAF1* in a micro-dissected dataset in patient tumor (red) and stroma (blue). (F) 3’-RACE of altered PDA-associated genes in MiaPaCa2 and Suit2 cells (representative images, n=3). Approximate length of the 3’-UTR form is denoted beside each band. (G) 3’-RACE of select genes in primary patient samples.

PDA samples are often characterized by substantial stromal contamination^5^; therefore, we sought to determine if significant APA events were present in the stroma or the tumor epithelium. First, we analyzed PDUI changes in a subset of 69 high purity TCGA-PAAD tumor samples^4^ (>33% tumor content). 89% of gene hits from our original analysis showed up as significant hits in the high purity dataset, suggesting that the majority of the detected APA changes were not attributable to stromal contamination (Supp. Fig. 2A,B). We further addressed this concern by visualizing the 3’-UTR profile of our candidate genes in an independent dataset containing RNA-seq information from 65 matched human PDA samples with micro-dissected tumor epithelia and stroma^5,60^. As an example, Fig. 1E shows the differential 3’-UTR shortening of *FLNA* and *PAF1* in patient tumor epithelium (tumor cells) as compared to the matched stroma.

We validated the presence of alternative 3’-UTR forms for several APA-regulated candidate genes by 3’-RACE (rapid amplification of 3’ ends) in 2 human pancreatic cancer cell lines (Suit2, MiaPaCa2) and 3 primary patient samples (Fig. 1F,G). These genes included the previously described PDA growth-promoting genes, as well as the spermine/spermidine acetyltransferase *SAT1*, and PP2A subunit B isoform δ (*PPP2R2D*). SAT1 modulates cell migration and resistance in multiple tumor types, while PPP2R2D is a component of the tumor suppressive phosphatase PP2A^61–66^. With the exception of *PPP2R2D*, which displayed significant 3’-UTR lengthening and downregulation in tumors, all of the validated genes were significantly shortened and overexpressed in the TCGA-PAAD dataset. We detected 3’-UTR short and long forms via 3’-RACE. The short 3’-UTR form for each shortened gene predominated over the long form (Fig. 1F,G). *ENO1* showed a single 3’-UTR form suggesting that this is the predominant form in cancer cells. In contrast, *PPP2R2D* showed an increased proportion of the 3’-UTR long form in PDA cell lines and patient samples as compared to the short form, suggesting greater use of the distal PAS for this putative tumor suppressive gene. For every candidate, we successfully identified PAS sites within its 3’-UTR sequence that matched the expected position of proximal and distal PAS in the detected 3’-RACE forms (Supp. Fig. 2C). Therefore, a large-scale comparison of 3’-UTR alterations can identify tumor epithelium-specific changes from the TCGA and GTEx datasets, and these 3’-UTR forms can be detected in cell models and patient samples.

### 3’-UTR changes are widespread among PDA patients and enriched in PDA pathways

To visualize the landscape of APA across PDA, we clustered patients (columns) based on change in PDUI score (tumor - normal mean; ΔPDUI) for 3’-UTR altered genes (rows) (Fig. 2A). This analysis uncovered a subset of genes (n=68) that showed 3’-UTR shortening (red) in >90% of patients, highlighting the widespread nature of APA across PDA. A smaller subset of 3’-UTRs (n=26, bottom heatmap) was recurrently lengthened (blue) in the tumor cohort. Hierarchical clustering identified multiple patient subgroups characterized by 3’-UTR alterations of specific gene sets (Subgroups 1-5). Notably, Subgroup 5 was enriched in shortened 3’-UTRs and contained relatively few lengthening events. In contrast, Subgroup 1 displayed fewer 3’-UTR shortening events and was enriched in 3’-UTR lengthening. Subgroups 2-4 were characterized by shortening events in specific subsets of genes. APA-based clustering therefore revealed distinct patient subgroups. These subgroups did not correlate with the mutational status of recurrently mutated PDA genes (*KRAS, CDKN2A, SMAD4, TP53*), nor did they associate with previously described PDA subtypes.

**Figure 2.**
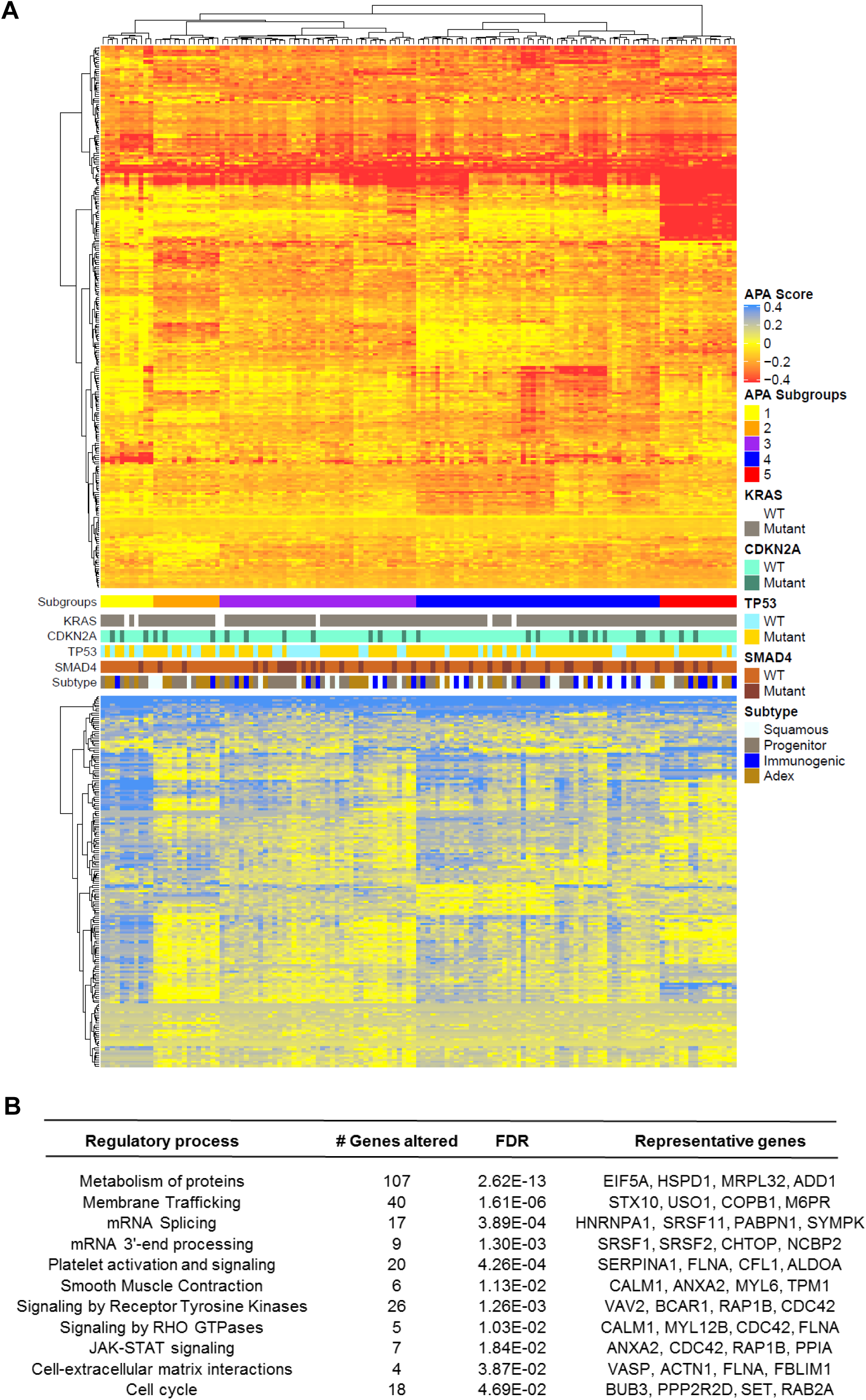
3’-UTR changes are widespread among PDA patients and enriched in PDA pathways. (A) The heatmap shows genes (rows) undergoing 3’-UTR shortening (red) or lengthening (blue) in each patient tumor (columns) compared to median score in normal pancreas for that gene. The profile of *KRAS, CDKN2A, TP53, SMAD4* mutations as well as tumor subtype is shown in the context of distinct APA-derived patient subgroups. (B) Significantly enriched (FDR<0.05) reactome pathways associated with 3’-UTR altered genes.

Pathway analysis of the significantly altered genes revealed enrichment for mRNA 3’ end processing and splicing, as well as smooth muscle contraction and platelet activation. Similar pathways have been found by pan-cancer APA analyses, concordant with the presence of recurrent APA events across multiple cancer types^41,43^. However, we observed further enrichments in PDA-associated pathways, including protein metabolism, signaling by receptor tyrosine kinases, signaling by RHO GTPases, JAK-STAT signaling and cell-extracellular matrix interactions (Fig. 2B). Therefore, APA alterations may regulate the activity of PDA-promoting pathways.

### 3’-UTR shortening identifies a poor prognostic cohort in PDA patients

Next, we asked whether APA events added additional prognostic information to PDA patient outcomes above the usual demographic and clinical factors: age, race, sex, stage, grade and surgical outcome. We selected genes with significant 3’-UTR alterations and univariate prognostic value, defining prognostic classes based on multivariate clustering (Fig. 3A). This segregated patients into three cohorts based on their 3’-UTR patterns (long=blue; short=red). Cohort A was predominantly associated with proximal PAS usage of genes from Groups 2 and 3, while cohort C was associated with distal PAS usage of the same genes. For Group 1 genes, distal PAS usage was predominant in cohort A while proximal PAS usage was predominant in cohort C. Neither patient cohort correlated with any of the known PDA tumor subtypes. Importantly, cohorts A and C displayed significant differences in overall survival, with patients in cohort C living significantly longer than those in cohort A (p=0.02) (Fig. 3B). Therefore, patterns of APA can be used as an independent prognostic indicator in PDA.

**Figure 3.**
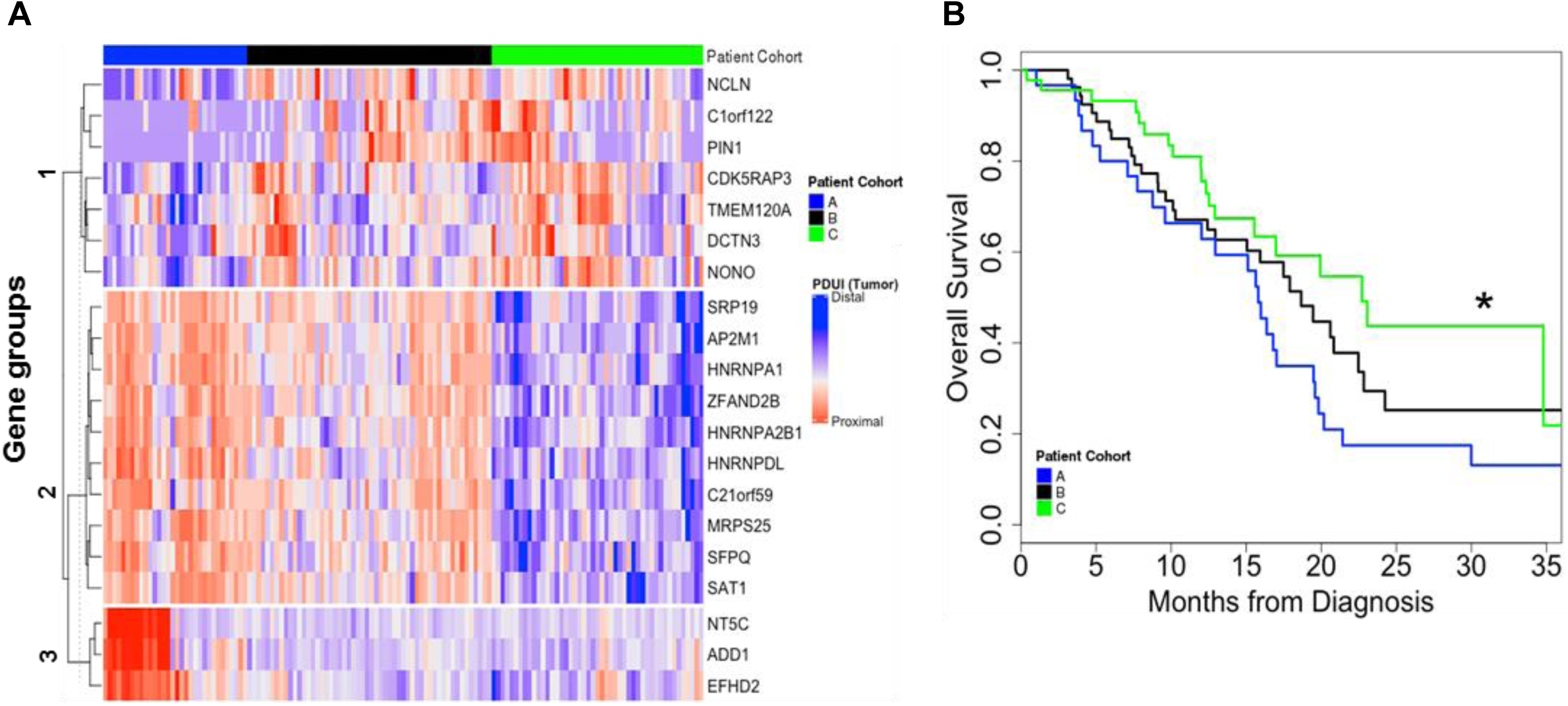
APA events identify a poor prognostic cohort in PDA patients. (A) Patients were clustered based on 3’-UTR short (red) and long forms (blue) of 3’-UTR altered genes (clustered into gene groups 1,2,3) and segregated into patient cohort A (blue), patient cohort B (black) and patient cohort C (green). (B) Kaplan-Meier survival plot for patient cohort A (blue), patient cohort B (black) and patient cohort C (green) (*p<0.05).

### Heterogeneity of proximal PAS usage of metabolic genes in PDA patients

Processes generating genetic and epigenetic heterogeneity can drive tumor evolution^67–69^. We hypothesized that APA could represent such a process, creating a diverse set of 3’-UTR forms and allowing cancer cells to select for those that promote their survival and propagation. To examine this heterogeneity, we compared the extent of proximal PAS usage across patients in any given gene between tumor and normal samples. *ALDOA* is shown as an example gene that exhibited a tight distribution of proximal PAS usage across normal as well as patient tumors (Fig. 4A). The left shift of the tumor sample mean score represents the expected shortening of the *ALDOA* 3’-UTR. However, for *FLNA*, while the normal samples had a tight distribution, PDA patients showed greater heterogeneity in proximal PAS usage (Fig. 4B). An analysis of heterogeneity in proximal PAS usage for all genes revealed that while the majority of genes did not show a significant change between normal and tumor conditions, 68 genes showed greater heterogeneity in tumor (orange) samples and only 9 genes showed greater heterogeneity in normal (purple) samples (Fig. 4C). This heterogeneity was not due to intrinsic differences between the TCGA and GTEx datasets because none of the 215 housekeeping genes in the dataset showed heterogeneity in the extent of proximal PAS usage^70,71^. The subset of 68 genes was enriched in metabolic genes, specifically amino acid transporters and purine metabolism. A wide range of heterogeneity of proximal PAS usage in PDA patients suggests a possible role of PAS usage plasticity in promoting cancer cell survival and progression.

**Figure 4.**
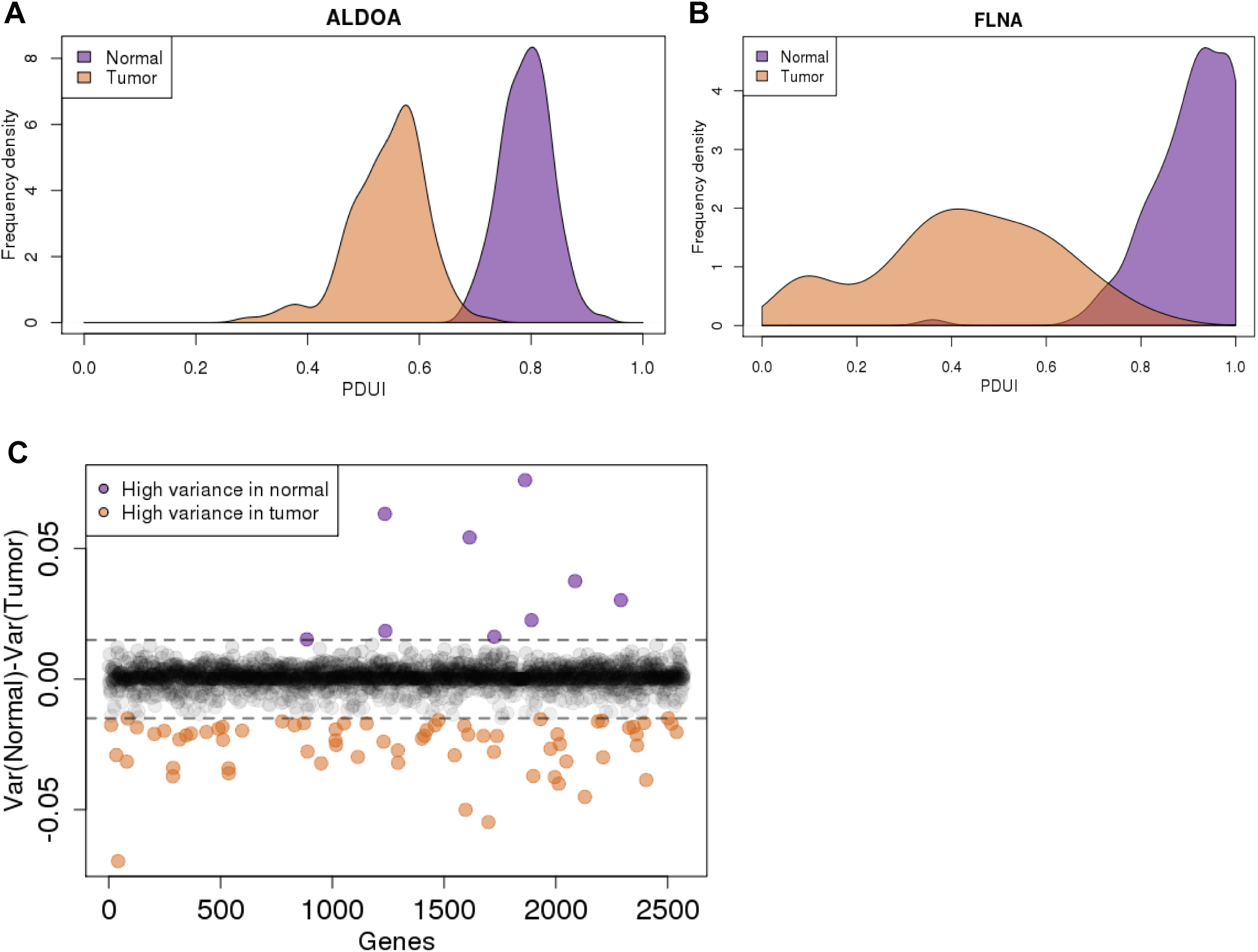
PDA patients show substantial heterogeneity in the extent of proximal PAS usage of metabolic genes. (A) Example of a 3’-UTR shortened gene (*ALDOA*) that has a tight distribution of its proximal PAS usage in normal pancreas (purple) as well as PDA patients (orange). (B) A 3’-UTR shortened gene (*FLNA*) that has a tight distribution in normal pancreas (purple); however, the extent of proximal PAS usage varies greatly across PDA patients (orange). (C) Plot of variance in PDUI for all genes between tumor and normal. Purple dots represent genes with high variance in normal samples while orange dots represent genes with high variance in tumor samples. Dashed lines represent 0.015 and –0.015 cutoffs.

### APA drives altered protein expression in PDA

To determine whether the identified APA events drive altered gene expression in PDA, we computed differential gene expression between normal (GTEx) and tumor (TCGA-PAAD) tissues. This allowed association studies between specific APA events and changes in gene expression. Among 3’-UTR shortened genes, 76 were significantly upregulated, while 50 genes were significantly downregulated in tumors (Fig. 5A,B). The pattern of 3’-UTR shortening preferentially associated with increased gene expression is consistent with pan-cancer APA analyses and conforms to the expectation that 3’-UTR shortened genes can escape miRNA regulation leading to increased gene expression^30,72,73^. In contrast, 3’-UTR lengthened genes showed a similar number of significantly upregulated (n=42) and significantly downregulated (n=41) genes, consistent with pan-cancer analyses, and most likely reflective of positive and negative regulation by RNA-binding proteins^29,74,75^.

**Figure 5.**
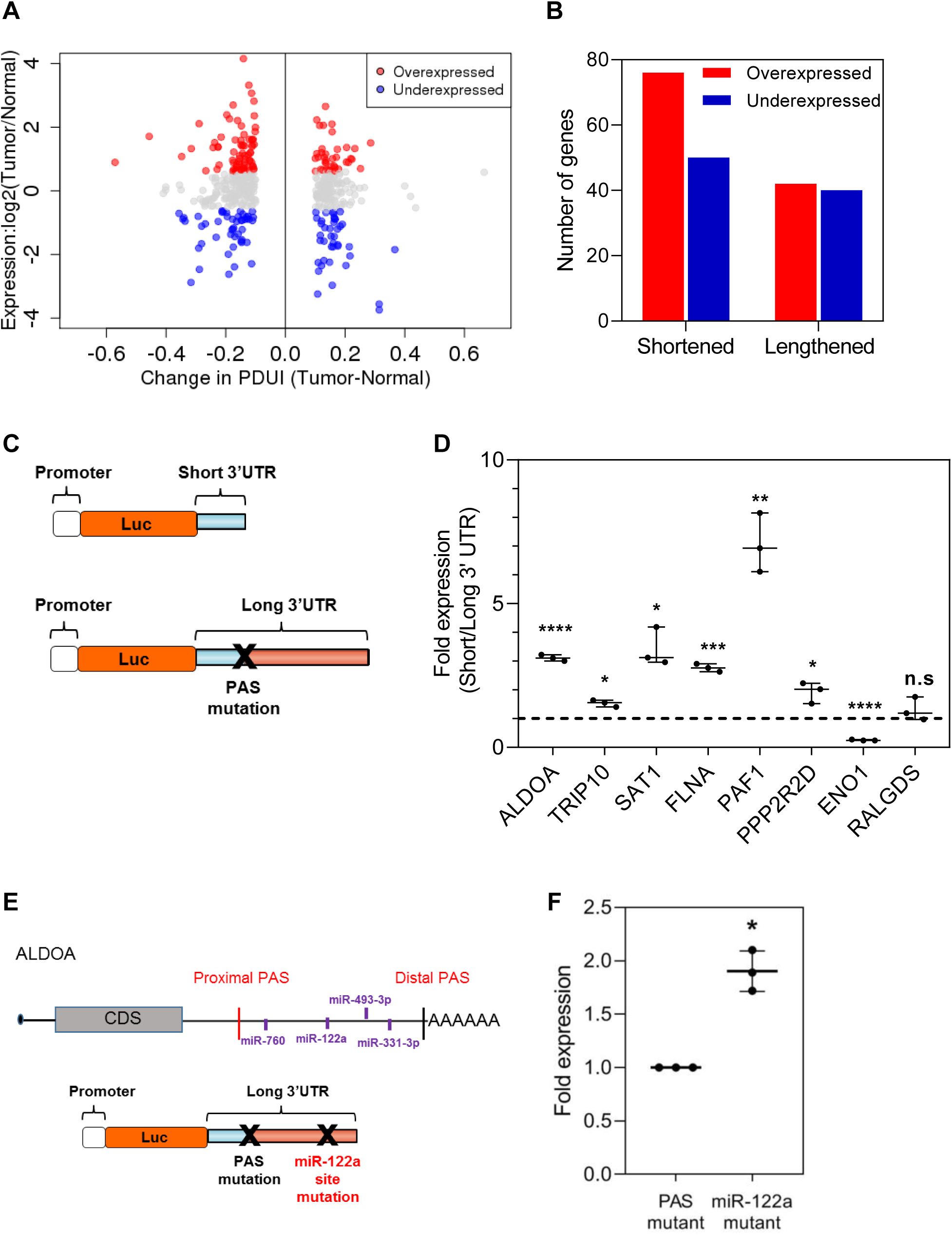
APA drives altered gene expression in PDA. (A) Log fold change in gene expression is plotted against ΔPDUI for 3’-UTR altered genes. Overexpressed genes (red dots) and underexpressed genes (blue dots) on the left represent 3’-UTR shortened hits while those to the right represent 3’-UTR lengthened hits. (B) Quantification of 3’-UTR altered genes that are overexpressed (red) or underexpressed (blue) in PDA tumors. (C) Schematic illustrating the luciferase reporter constructs. (D) Normalized fold expression change of the luciferase reporter (short 3’-UTRs / long-3’UTRs) for the selected list of 3’-UTR altered genes (n=3). The long 3’-UTR expression for each gene is normalized to 1. Each whisker plot denotes the median as the center line and the minimum and maximum values as the whiskers (*p<0.05, **p<0.01,***p<0.005, ****p<0.001). (E) Schematic showing the *ALDOA* 3’-UTR with positions of conserved miRNA sites as well as the miRNA mutant construct used. (F) Fold expression change of miRNA mutant construct compared to the PAS mutant in luciferase assays (n=3). The PAS mutant expression is normalized to 1.

To experimentally validate the relationship between APA and protein expression, we performed luciferase reporter assays in MiaPaCa2 cells, comparing protein expression driven by short and long 3’-UTRs (Fig. 5C). We focused on the candidate oncogenes and tumor suppressors validated by 3’-RACE and that showed significant association between 3’-UTR changes and gene expression in tumors. These candidates included *ALDOA, FLNA, PAF1, TRIP10, ENO1*, *SAT1* (shortened and upregulated in tumors) and *PPP2R2D* (lengthened and downregulated in tumors). We also included *RALGDS* which is shortened but does not show altered expression in tumors. We cloned the short and long 3’-UTRs of each gene (estimated via 3’-RACE) downstream of a *Renilla* luciferase reporter and measured luminescence as a readout of protein expression (Fig. 5C). To ensure that the long 3’-UTR form for each reporter gene remained intact (*i.e.*, did not undergo APA-mediated shortening upon transfection into cells), we mutated their functional proximal PAS. For all genes tested except *ENO1* and *RALGDS*, the short 3’-UTR form showed significantly increased luminescence compared to the long 3’-UTR form (Fig. 5D). As predicted, the 3’-UTR short and long forms of *RALGDS* showed similar expression. In contrast to our expectations, the short form of *ENO1* showed decreased protein expression suggesting that 3’-UTR shortening is not the sole mechanism regulating protein abundance of *ENO1* in PDA. These results also reinforce the observation that shorter 3’-UTRs do not always increase protein expression^30^. Overall, the above results suggest that APA-mediated 3’-UTR alterations can regulate the protein expression of growth-promoting genes in PDA cells.

We next sought to determine the mechanism underlying the 3’-UTR-mediated gene regulation of the PDA oncogene *ALDOA*. Given that miRNAs primarily destabilize their target mRNA and that *ALDOA* undergoes 3’-UTR shortening and upregulation, we searched the *ALDOA* 3’-UTR for putative miRNA binding sites that would be lost upon PDA-associated shortening (Fig. 5E). We identified the tumor suppressive miRNA miR-122a within this lost region; miR-122a is highly expressed in PDA cell lines^76,77^. Mutation of the miR-122a site within the long 3’-UTR of *ALDOA* significantly restored protein expression (Fig. 5F). Therefore, altered APA can regulate oncogene expression in PDA through modulation of available regulatory miRNA binding sites.

### APA-mediated loss of tumor suppressive miRNA binding sites is associated with poor patient outcome

To assess global patterns of APA-mediated miRNA binding site loss we searched for highly conserved miRNA binding sites (conserved across human, mouse, rat, dog and chicken) within the lost 3’-UTRs of all shortened genes. This analysis revealed that 42% of genes lost at least one highly conserved miRNA binding site (Fig. 6A), suggesting that alteration of the miRNA binding site repertoire is an important mode of APA-mediated regulation. Next, we sought to determine if any miRNA families were preferentially lost in shortened 3’-UTRs of PDA patients. We computed an index for repression for each miRNA family as a function of the miRNA site context scores (obtained from TargetScan) and the abundance of the 3’-UTR form containing that site. This index was then compared between PDA patients and normal controls to yield a Z-score. A lower Z-score for a miRNA family reflects preferential loss of its binding sites due to 3’-UTR shortening. Interestingly, 6 of the top 8 identified miRNAs have been implicated as tumor suppressors in PDA, including miR-329 and miR-133a^78–82^ (Fig. 6B). These results suggest that APA regulates oncogenic gene expression through preferential loss of tumor suppressive miRNA binding sites and may therefore confer a selective advantage to the cell.

**Figure 6.**
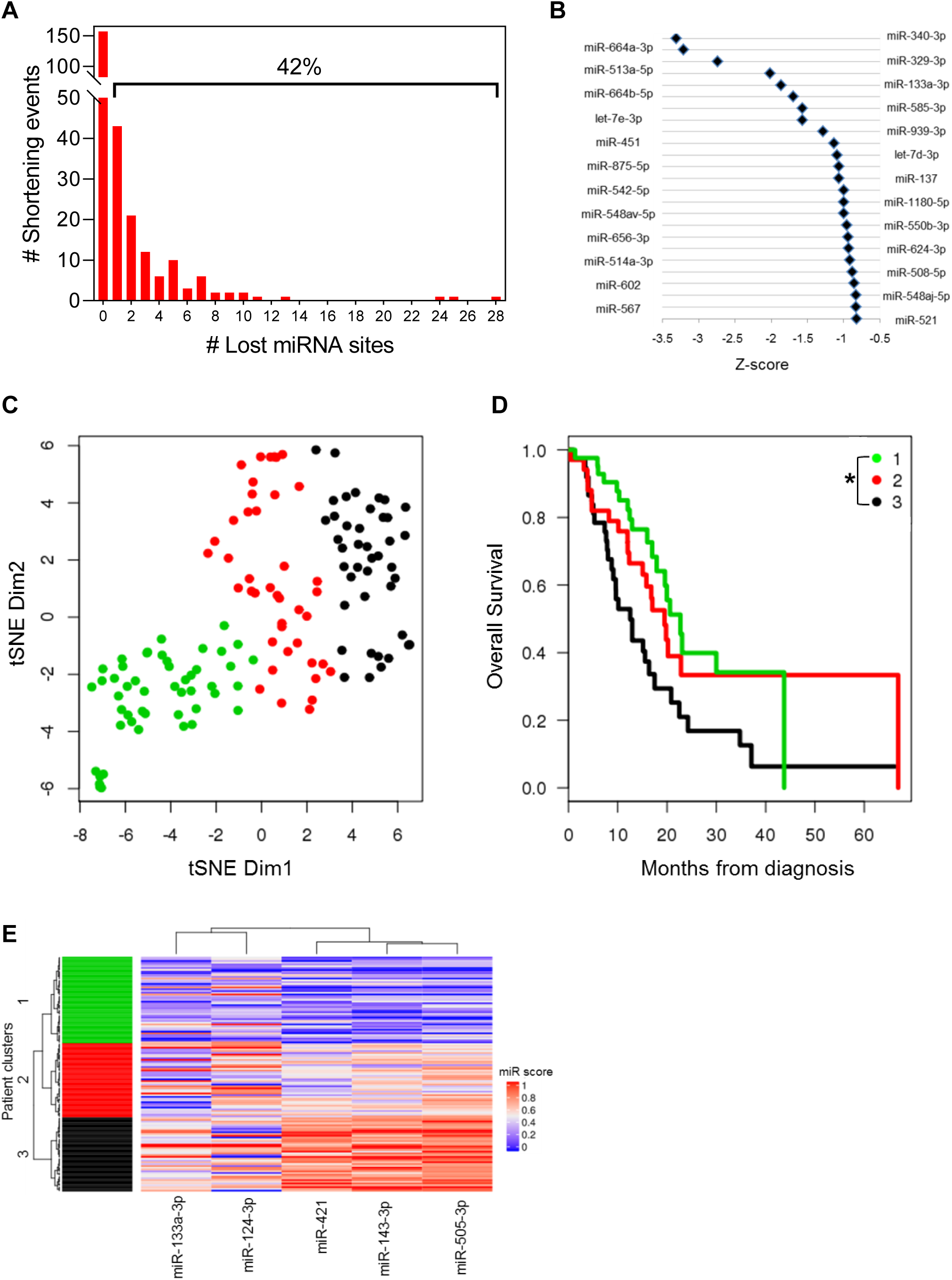
APA-mediated loss of tumor suppressive miRNA binding sites is associated with poor patient outcome. (A) Number of genes that lose highly conserved miRNA binding sites due to 3’-UTR shortening. The percentage of genes that lose at least 1 miRNA binding site is indicated above the bracket. (B) Highly conserved miRNA families were plotted against their Z-score, an index of the lost binding sites where a more negative Z-score indicates more significant binding site loss. (C) tSNE plot depicting TCGA patient clusters in the highly conserved miRNA feature space. (D) Kaplan-Meier survival plot for the 3 patient clusters identified in (C) (*p<0.05 for Cluster 1 to Cluster 3 comparison). (E) Heatmap depicting the association of miRNA binding site loss (miR score) with patient clusters.

Next, we determined whether loss of specific miRNA sites associated with 3’-UTR alterations is associated with patient outcome. We quantified loss of highly conserved miRNA binding sites for each patient as a function of the extent of proximal PAS usage in all genes that lost those miRNA sites (see Methods). Clustering in the miRNA feature space revealed 3 patient groups (Fig. 6C) with significant differences in overall survival (p=0.012 between Clusters 1 and 3; Fig. 6D). The miRNAs most significantly associated with the patient clusters included miR-133a, miR-124, miR-421, miR-143 and miR-505. Binding sites for each miRNA were preferentially lost from Cluster 1 as compared with Cluster 3, suggesting that loss of these regulatory sites correlates with poor survival of PDA patients (Fig. 6E). Indeed, miR-133a, miR-124 and miR-143 are known tumor suppressors in PDA, again supporting the role of APA in selective loss of tumor suppressive miRNA binding sites^80,83–89^.

### The APA-regulated gene *CSNK1A1* is required for proliferation and clonogenic growth of PDA cells

Our data showed APA-mediated regulatory changes in genes known to promote PDA pathogenesis. We hypothesized that our altered gene list may also contain growth-promoting genes not previously implicated in PDA biology, and therefore new therapeutic targets. We focused on the subset of druggable genes that were significantly shortened and upregulated in PDA. Finally, we overlaid this list with results from a genome-wide CRISPR screen, identifying genes essential for PDA cell proliferation^90^. This analysis identified *CSNK1A1*, the gene encoding the serine/threonine kinase casein kinase 1α (CK1α). CK1α regulates the Wnt/β-catenin signaling pathway and has dual functions in cell cycle progression and cell division^91–93^. CK1α is known to influence tumor progression; however, its role as a tumor suppressor or oncogene is tumor type-dependent^91,93–95^ and CK1α has no known roles in PDA. *CSNK1A1* has very low gene expression in normal pancreas but is overexpressed in PDA^96^. We found that *CSNK1A1* shows significantly higher expression in the PDA epithelium as compared to precursor lesions (premalignant pancreatic intraepithelial neoplasia (PanIN) (Fig. 7A) and intraductal papillary mucinous neoplasia (IPMN)). We found no significant difference in *CSNK1A1* expression in the stroma between PDA and precursor lesions. We then determined whether differential CK1α activity mediates progression from precursor lesions to PDA. As a first step, we assembled a context-specific gene regulatory network from 242 micro-dissected epithelial gene expression profiles using the Algorithm for Reconstruction of Accurate Cellular Networks (ARACNe)^97,98^. The input list of regulatory proteins for ARACNe contained DNA binding domain containing proteins as well as signaling proteins (including CK1α) and therefore, was not restricted to transcription factors alone. We then employed MARINa (MAster Regulator INference algorithm) to determine the activity of CK1α between precursor lesions and PDA samples as a function of expression of the CK1α regulon (inferred using ARACNe)^99^. If the CK1α targets are enriched for genes that are differentially expressed between precursor lesions and PDA, it indicates differential CK1α activity between the two conditions. Indeed, the positive targets of CK1α were more highly expressed in PDA epithelium, whereas the negative targets showed increased expression in precursor lesions, indicating that CK1α activity may promote the progression from precursor lesions to PDA (Fig. 7B). Importantly, CK1α differential activity was not present in the stroma between PDA and precursor samples suggesting the specific role of CK1α in PDA epithelium. As predicted by our computational analysis, 3’-RACE showed that *CSNK1A1* has an increased proportion of the short 3’-UTR form as compared to the long 3’-UTR form in PDA cells (Supp. Fig. 3A).

**Figure 7.**
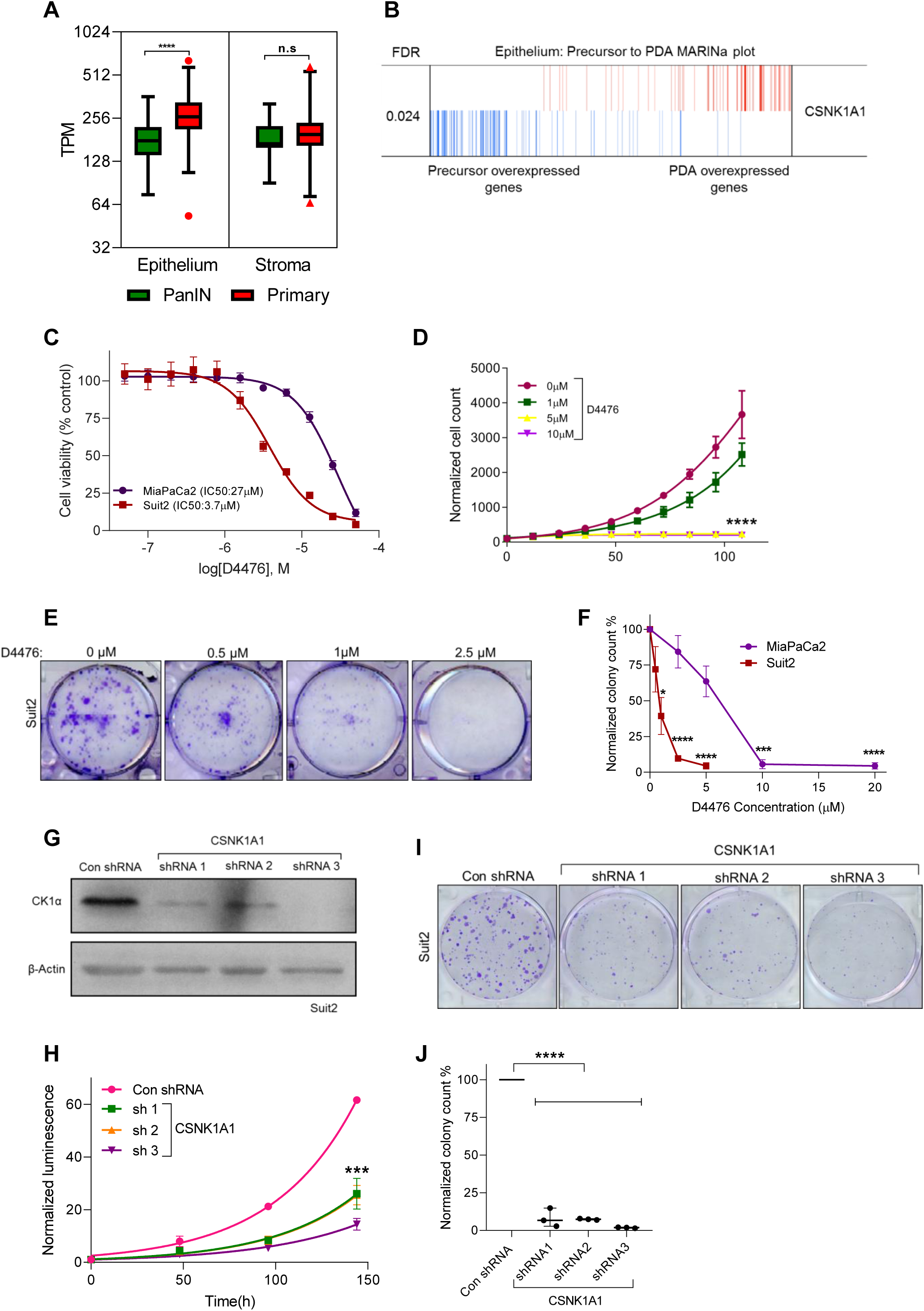
CK1α is required for cell proliferation and is a putative drug target in PDA. (A) A plot showing *CSNK1A1* gene expression (in transcripts per million) in PDA (red) as compared to PanIN lesions (green) in the epithelium and stroma from micro-dissected samples (****p<0.001). (B) MARINa plot showing CK1α targets (blue: negative targets, red: positive targets) ranked by their differential gene expression from precursors (left) to PDA epithelium (right). (C) Dose-response of MiaPaCa2 (purple) and Suit2 (red) cell lines to the CK1α small molecule inhibitor, D4476 (n=3). (D) Cell proliferation of Suit2 cells treated with indicated doses of D4476 (n=3, ****p<0.001). (E) Clonogenic growth assay of Suit2 cells treated with indicated drug doses. (F) Quantification shows the number of colonies in (E) (n=3, ***p<0.005, ****p<0.001). (G) A representative blot confirming CK1α knockdown in Suit2 cells with a non-targeting control shRNA (Con shRNA) or with one of three different shRNAs targeting *CSNK1A1* (n=3). (H) Cell proliferation of Suit2 control and CK1α knockdown cells (n=3, ***p<0.005). (I) Clonogenic growth assay of control and CK1α knockdown cells (n=3). (J) Quantification shows the number of colonies in (I) (****p<0.001).

We then investigated the potential for CK1α inhibition to regulate PDA biology with the widely used small molecule inhibitor D4476^94,96,100^. We treated the PDA cell lines MiaPaCa2 and Suit2 with D4476; while MiaPaCa2 and Suit2 cells were both sensitive to D4476 treatment, Suit2 cells displayed a 10-fold lower IC50 (Fig. 7C). Both cell lines also showed dose-dependent decreases in cell proliferation (Fig. 7D, Supp. Fig. 3B) and clonogenic growth in the presence of the inhibitor (Fig. 7E,F, Supp. Fig. 3C). To provide genetic evidence for the role of CK1α in PDA cell growth, we knocked down *CSNK1A1* in Suit2 and MiaPaCa2 cells with 3 short hairpin RNAs (shRNA) (Fig. 7G, Supp. Fig. 3D). In concordance with the pharmacological results, *CSNK1A1* knockdown decreased both cell proliferation and clonogenic growth of PDA cells (Fig. 7H-J, Supp. Figs. 3E,F), with Suit2 cells showing increased sensitivity to CK1α loss. The strongest phenotypic effects were associated with the most efficient knockdown (shRNA 3) in both cell lines. Therefore, we identify CK1α as a putative drug target in PDA and reveal the potential of cancer-specific APA analyses to identify mechanisms of altered gene expression driving cancer pathogenesis.

## DISCUSSION

Dysregulated gene expression is a cardinal feature of cancer^101^. However, how gene expression is altered in cancer and whether the processes driving this dysregulation can be targeted therapeutically are areas of active investigation. APA has recently been identified as a candidate driver of gene expression dysregulation. APA factors frequently show aberrant expression in cancer, modulate the expression of known oncogenes and tumor suppressors, and knockdown studies have highlighted their relevance to the cancer phenotype^31–34,102,103^. Whole-genome CRISPR and shRNA screens have also revealed the requirement for several APA factors in pancreatic cancer cell growth (www.depmap.org). Global analyses have revealed widespread 3’-UTR changes across multiple cancer types, uncovering recurrent alterations common across the cancer spectrum^38–41^. Recent findings suggest that while some APA events are widely shared across cancers, many are tumor type-specific^42^. Despite this observation, there have been few attempts to study APA in a single tumor type with sufficient power to identify tumor-specific alterations and vulnerabilities.

To our knowledge, this study represents the first global, in-depth, single cancer view of APA, and the first examination of APA in PDA clinical samples. The only previous study of APA in PDA showed gemcitabine-induced 3’-UTR shortening of the transcription factor ZEB1 in the context of drug resistance^104^. Previous APA analyses combined multiple tumor types and used tumor-adjacent tissue as a “normal” control. However, matched tumor-adjacent normal tissues are known to represent a state that significantly differs from healthy, normal tissues and may therefore miss critical APA events^48^. Furthermore, there are insufficient numbers of tumor-adjacent pancreatic samples within TCGA for a statistically stringent analysis. Therefore, we attempted to address these issues by using normal pancreas RNA-seq information from the GTEx project. An important limitation of comparing independently collected datasets is the inherent disparity between them. We attempted to rectify this by: a) confirming that the two datasets underwent identical library preparation methods on the same type of sequencing platform; b) following identical data processing pipelines from the raw sequencing data to generate the coverage data; c) validating our top hits in an independent micro-dissected dataset. Consistent with previous publications comparing TCGA and GTEx datasets, we observed minimal batch effects. As batch effects cannot be completely ruled out, we performed experimental validation of several candidate APA regulated genes, including *PAF1* and *ALDOA*, highlighting the robustness of our approach and relevance of our findings to PDA biology. Furthermore, this approach will allow the analysis of APA in other tumor types for which little tumor-adjacent material is present in TCGA.

Multiple insights from our analyses are noteworthy. We find that APA events are recurrent and widespread across PDA patients. For example, 68 genes were shortened and 28 genes were lengthened in greater than 90% of the patient cohort. This supports the conjecture that APA dysregulation is a frequent event in PDA. In support of this hypothesis, we find that several APA factors are highly expressed in PDA, including *CSTF2* (Supp. Fig. 4). CSTF2 has previously been implicated as a promoter of lung and bladder cancer, through the regulation of *ERBB2* and *RAC1* 3’-UTRs, respectively^33,34^. We find frequent 3’-UTR alterations in several notable PDA-relevant genes whose mechanisms of regulation were previously unknown, including *PAF1*, *ALDOA* and *FLNA*. Many of the shortened 3’-UTRs correlated with increased gene expression, providing the first collection of 3’-UTR alterations that correlate with gene expression changes in PDA. We were able to functionally validate these through luciferase reporter assays, highlighting the robustness of our analysis. Consistent with pan-cancer APA analyses, we find enrichment for pathways such as smooth muscle contraction and mRNA 3’-end processing^29,41,43^. However, we also find enrichment for pathways and processes implicated in PDA biology, including protein metabolism, receptor tyrosine kinase signaling and signaling by RHO GTPases. Indeed, the link between 3’-UTR alterations and cancer metabolism has been identified in previous pan-cancer APA analyses^41^. We also find an unexpected enrichment for loss of binding sites for tumor-suppressive miRNAs in frequently lost 3’-UTR regions. Therefore, we propose that APA is an underappreciated mechanism of gene dysregulation in PDA, driving the expression of growth-promoting genes through disruption of miRNA-mediated regulation.

The extent of heterogeneity in proximal PAS usage across cancer patients has been largely overlooked in previous pan-cancer APA analyses. We found little heterogeneity in the extent of 3’-UTR proximal site usage in most genes (including housekeeping genes) in both normal and PDA samples, again providing evidence for minimal batch effects. However, PDA patients showed substantial heterogeneity in the extent to which their metabolic genes used the proximal PAS. This metabolic plasticity in turn could serve as a mechanism to deal with the fluctuating metabolic demands of cancer cells. These data support the possibility that APA may drive deregulation of cancer metabolism and tumor evolution by allowing for PAS choice plasticity of critical metabolic genes in PDA.

Several studies have demonstrated the power of APA analysis to improve expression-based prognostic markers. We report the first subset of 3’-UTR alterations that act as an independent prognostic indicator of PDA outcome. While several of the genes in this set are known regulators of tumorigenesis, including *SAT1*, many have not been implicated in PDA biology and may represent new mediators of cancer phenotypes. Interestingly, lost miRNA sites are enriched for tumor-suppressive miRNA families. In particular, we observed that patients who retain binding sites for a subset of 5 miRNAs survive longer than patients who lose them. This uncovers the prognostic role for a novel subset of miRNA mediators in PDA.

Our in-depth analysis of APA in PDA revealed a critical role for the druggable target CK1α in PDA cell growth and survival. While CK1α has known roles in Wnt signaling and p53 activation, important mediators of PDA progression, the relevance of CK1α to PDA was previously unknown^93–96^. Furthermore, the mechanisms of regulation of CK1α in cancer are not well understood, although promoter methylation in melanoma has been reported^105^. Interestingly, two CK1α isoforms have been detected in HeLa cells, with the shorter isoform being generated from the use of an alternative PAS^106^. We show that CK1α exhibits increased activity in PDA samples as compared to precursors, and that pharmacological and genetic blockade of CK1α attenuates PDA cell proliferation and clonogenic growth. Therefore, our single-cancer approach can identify APA-regulated, disease-specific vulnerabilities.

Our computational analysis and experimental validation have revealed unexpected mediators of PDA biology and broadened our understanding of the regulatory role of 3’-UTR sequence space in cancer. This comprehensive analysis reveals the scope of previously uncharacterized APA events in regulating functionally relevant PDA genes, improving patient prognosis and driving tumor evolution. We propose that the landscape of 3’-UTR alterations in PDA represents a novel avenue to better understand PDA progression and identify new drug targets.

## DATA AVAILABILITY STATEMENT

All RNA-seq files were downloaded via NCBI dbGAP. This included 184 normal pancreas SRA files from GTEx (dbGAP accession phs000424.v8.p2) and 148 BAM files within the TCGA-PAAD cohort (https://portal.gdc.cancer.gov/).

## ACKNOWLEDGEMENTS

This work was supported by NCI grants P30 CA016056 and R25 CA181003, an award from the Roswell Park Alliance Foundation to MEF, and DoD grant OC170368 to KHE. We thank the Roswell Park Gene Modulation core, Pathology Shared Resource, Genomics Shared Resource and the Small Molecule Screening Shared Resource for their assistance. We thank the members of the Feigin Lab, the Roswell Park Department of Pharmacology and Therapeutics, and the Science Twitter community for their helpful comments on the manuscript.

## AUTHOR CONTRIBUTIONS

Wrote the manuscript: SV, MEF

Supervised the study: MEF

Performed DaPars analysis: SV

Performed biological experiments: SV, AAT, JRS, AAA

Contributed to data analysis: SV, KHE, HCM, KPO

Developed prognostic signatures: KHE

## Supplementary Figure Legends

**Supplementary Figure 1. Analysis flowchart.** We identically processed raw RNA-seq data from the GTEx and TCGA-PAAD cohorts to analyze APA events in PDA. Predicted genes were further validated using a smaller high purity TCGA cohort and an independent micro-dissected dataset. The resulting genes were interrogated for associated APA trends, prognostic significance and gene expression changes.

**Supplementary Figure 2. Gene hits in the high purity TCGA-PAAD subset.** (A) Volcano plot depicting significant gene hits (FDR<0.01) whose |ΔPDUI| > 0.1 in the 69 high purity samples (> 33% tumor content). (B) Venn diagram representing the overlap in significant gene hits between the DaPars analysis of 148 TCGA-PAAD samples and the 69 high purity TCGA-PAAD dataset. (C) 3’-UTR schematic and sequence of 3 example candidate genes (*FLNA, PPP2R2D* and *PAF1*). The stop codon is highlighted in blue and marks the beginning of the 3’-UTR sequence. The functional PAS sites estimated from 3’-RACE forms are highlighted in red.

**Supplementary Figure 3. CK1α is required for cell proliferation and is a putative drug target in PDA.** (A) 3’ RACE of *CSNK1A1* in Suit2 and MiaPaCa2 cells (representative images from 3 independent experiments). (B) Cell proliferation of MiaPaCa2 cells treated with indicated doses of D4476 (n=3, ****p<0.001). (C) Clonogenic growth assay of MiaPaCa2 cells treated with indicated drug doses. (D) A representative blot (n=3) confirming CK1α knockdown in MiaPaCa2 cells with a non-targeting control shRNA (Con shRNA) or one of three different shRNAs targeting *CSNK1A1*. (E) Cell proliferation of MiaPaCa2 control and CK1α knockdown cells (n=3, ***p<0.005). (F) Clonogenic growth assay of MiaPaCa2 control and CK1α knockdown cells (n=3).

**Supplementary Figure 4. APA factor expression in PDA.** Fold expression change of core APA factors between TCGA (tumor) and GTEx (normal pancreas). Dotted lines represent 1.5-fold (red) and 0.66-fold (blue) cutoffs.

## MATERIALS AND METHODS

### Data collection and preprocessing

Our study focused on PDA tumors consistent with the histology of PDA (n=148). All RNA-seq files were downloaded via NCBI dbGAP. This included 184 normal pancreata SRA files from GTEx (dbGAP accession phs000424.v8.p2) and 148 BAM files within the TCGA-PAAD cohort (https://portal.gdc.cancer.gov/). GTEx SRA files were aligned exactly according to the TCGA RNA-seq alignment pipeline using GENCODE.v22 annotations. Bedgraph files were generated using bedtoolsv2.26 and were supplied as input to the DaPars algorithm.

### DaPars analysis

DaPars processes bedgraph coverage files to identify differences in 3’-UTR lengths between two conditions. The output of our analysis contained putative 3’-UTR altered transcripts and was comprised of 2573 unique genes. The subset of genes that were significantly altered in their PDUI scores were calculated using Fisher’s exact test (|ΔPDUI| >0.1, FDR<0.05) between normal and PDA tumors. A similar analysis was performed with a subset of 69 high purity PDA tumor samples.

### Bioinformatics analyses and statistical methods

#### Analysis of heterogeneity

The variances in proximal PAS usage across tumor samples (Var(Tumor)) as well as normal samples (Var(Normal)) were computed for each gene and the difference (Var(Normal)-Var(Tumor)) was plotted (R version 3.5.2).

#### Heatmap analysis

A heatmap representing the extent of 3’-UTR alterations across PDA patients was generated (R version 3.4.3). For each significant gene hit (row), the median GTEx PDUI score was subtracted from the PDUI score for each TCGA PDA patient to obtain a measure of ΔPDUI (change in 3’-UTR length for that gene for each patient). Hierarchical clustering of patients (columns) segregated them into 5 distinct subgroups. Rows were similarly clustered to yield subsets of genes undergoing a higher degree of 3’-UTR shortening (red) or lengthening (blue). The mutational status of commonly altered PDA genes and PDA subtype for each TCGA patient was highlighted.

#### Pathway analysis

PANTHER (Protein ANalysis Through Evolutionary Relationships) was used for pathway analysis^107,108^. The statistical overrepresentation test was used to statistically determine over or under-representation of reactome pathways in comparison to the reference list (all human genes in the PANTHER database) using Fisher’s exact test (FDR <0.05).

#### Survival analysis

We selected genes with significant 3’-UTR shortening for multivariate survival time model building by first computing the residuals from a multivariate proportional hazards model fit to clinical factors (age, stage, grade, surgical outcome, race and sex) and selecting only those genes with significant univariate correlation with this clinically unexplainable prognostic signal. We then used K-means clustering among selected genes to define 3 prognosis groups based on the within/between sum of squares criterion. The prognostic value of this classification is described by standard Kaplan-Meier plot and the log-rank test.

#### Differential gene expression analysis

Differential gene expression analysis between TCGA-PAAD and GTEx normal pancreas samples was performed using DESeq2. Genes showing (1) Fold change > 1.5 (2) FDR<0.05 (3) log_2_CPM > 3 were considered differentially expressed. The association between PDUI score and gene expression was plotted in R version 3.4.3.

#### Percentage of lost miRNA sites

Highly conserved miRNA binding sites and their genomic positions were downloaded from TargetScanHuman 7.2. This list, along with DaPars prediction of genomic coordinates of lost 3’-UTRs was used to plot the number of genes that lose at least 1 highly conserved miRNA binding site.

#### miRNA families preferentially associated with lost sites

In order to determine miRNAs associated with sites enriched in lost 3’-UTRs, miRNA target predictions and the cumulative weighted context++ scores (CWCS) were downloaded from TargetScanHuman 7.2. CWCS estimates the predicted cumulative repression for a miRNA at the site. The lost miRNA binding sites in the shortened 3’-UTRs of PDA patients were inferred from DaPars predictions. A weighted target site score was computed as the sum over all genes with shortened 3’-UTRs in tumor, with the CWCS of each target site for the miRNA multiplied by the normalized abundance of the gene’s 3’-UTR form in which the predicted target site was present. The fold-change (f) of the sum of weighted target site scores in lost 3’-UTR regions for PDA tumor over normal was calculated. The labels of the miRNA target sites were permuted to assess the significance of the fold-change. 1000 such randomizations were performed and the mean (m) and standard deviation (s) of the fold changes across the randomized data sets was computed. The significance of the fold change was computed in form of the Z-score defined as (f-m)/s. A lower Z-score indicates that the loss in miRNA binding sites is higher than that expected by chance.

#### miRNA prognostic signature

We quantified the impact of APA-based loss of miRNA binding as follows:

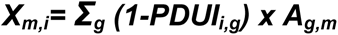

where ***A_g,m_*** is an indicator function that the short versus long 3’-UTR of the gene ***g*** contains the binding site for miRNA ***m***, the impact to the ***i^th^*** person is ***X_m,i_***. We used Sure Independence Screening (SIS) to search through all affected miRNAs and identify features that were associated with survival univariately^109^. To study the multivariate effect, we reorganized cases using the euclidean distance between SIS selected features, visualized with tSNE, and defined clusters with model-based Gaussian clustering using the BIC criterion to select cluster number. Survival differences were tested across all groups by the log-rank test and were visualized by Kaplan-Meier estimate. The pattern of loss of miRNA binding sites across patient clusters were visualized for a subset of miRNAs in a heatmap.

#### MARINa plot

The pancreatic cancer regulatory network was reverse engineered by ARACNe-AP from 242 microdissected epithelial gene expression profiles which were generated from 197 primary PDA, 26 low-grade PanIN and 19 low-grade IPMN lesions^5,60,98^. Raw counts were normalized to account for different library sizes after filtering out genes with less than one fragment per million mapped fragments in at least 20% of the samples, and the variance was stabilized by fitting the dispersion to a negative binomial distribution as implemented in the DESeq2 R package^110^. ARACNe was run with standard settings (using data processing inequality (DPI), with 100 bootstrap iterations using all gene symbols mapping to a set of 1856 transcription factors that includes genes annotated in the Gene Ontology (GO) molecular function database as GO:0003700 (‘transcription factor activity’), GO:0004677 (‘DNA binding’), GO:0030528 (‘transcription regulator activity’) or as GO:0004677/GO: 0045449 (‘regulation of transcription’), 671 transcriptional cofactors (a manually curated list, not overlapping with the transcription factor list, built upon genes annotated as GO:0003712, ‘transcription cofactor activity’, or GO:0030528 or GO:0045449) or 3,540 signaling pathway related genes (annotated in GO Biological Process database as GO:0007165 ‘signal transduction’ and in GO cellular component database as GO:0005622, ‘intracellular’, or GO:0005886, ‘plasma membrane’) as candidate regulators^111,112^. Thresholds for the tolerated DPI and mutual information P value were set to 0 and 10–8, respectively. For master regulatory analysis, we tested the differential activity for CK1α between precursor lesions and PDA by applying the multi-sample version of the VIPER algorithm (msVIPER)^99^. msVIPER considers the distribution of negative and positive targets of CK1α in the progression gene expression signature to infer its activity.

### Experimental methods

#### Cell lines, antibodies and general reagents

MiaPaCa2 and HEK293 cells were purchased from ATCC and cultured in DMEM media (Cat# MT 10-013-CV, Corning) and 10% fetal bovine serum. Suit2 cells were obtained from Dr. David Tuveson (Cold Spring Harbor Laboratory). Cell lines were periodically verified to be mycoplasma free using the Mycoalert kit (Cat# LT07-701, Lonza). All transfections were carried out using Lipofectamine 3000 (Cat# L3000008, Thermo Fisher Scientific) as per manufacturers protocol. All primers used in this study were purchased from Integrated DNA Technologies (IDT) and PCR reactions were performed using Q5 Hot start DNA polymerase (Cat# M0493L, NEB). cDNA synthesis was carried out using Superscript II Reverse Transcriptase (Cat# 18064022, Thermo Fisher Scientific). miRNA site mutations in *ALDOA* 3’-UTR as well as mutations at the proximal PAS of long 3’-UTRs were introduced using NEB Builder HiFi DNA assembly (Cat# E2621S, NEB). The *Renilla* reporter plasmid pIS1 (Plasmid# 12179) as well as the firefly plasmid pIS0 (Plasmid# 12178) were purchased from Addgene. Luciferase assays were performed using Dual Luciferase Reporter Assay System (Cat# E1910, Promega). For *CSNK1A1* drug studies, the small molecule inhibitor D4476 (Cat# 13305, Cayman Chemical) was dissolved in DMSO (Cat# S1078, Selleckchem) at a stock concentration of 20mM. For dose-response measurements and certain cell proliferation experiments, cell viability was assessed using CellTiter-Glo (Cat# G7571, Promega). 3 distinct predesigned shRNAs (sh1:Cat# V2LHS_176052, sh2:Cat# V2LHS_221905, sh3:Cat# V2LHS_263361) against *CSNK1A1* were procured from a commercial shRNA library (Dharmacon) from the Roswell Park Gene Modulation core. Primary antibodies used in this study included a polyclonal antibody against CK1α (Cat# A301-991A-M, Bethyl labs) and a monoclonal antibody against β-actin (Cat# 3700S, Cell Signaling Technology). Secondary antibodies included horseradish peroxidase-conjugated goat anti-mouse (Cat# A4416, Sigma) and goat anti-rabbit (Cat# 45-000-682, Fisher Scientific) antibodies.

#### Cell lysis and RNA extraction

MiaPaCa2 and Suit2 cells were grown to 100% confluence in 10cm plates. The cells were washed with 10mL PBS, and 1mL TRIzol was added to the cell culture plate. Cells were scraped, then incubated in a 1.5mL microcentrifuge tube for 5 minutes. 0.2mL of chloroform was added, mixed well and the tubes were incubated at room temperature for 2-3 minutes. The samples were centrifuged at 12000xg for 15 minutes at 4°C and the upper aqueous phase was transferred to a fresh tube. After addition and incubation with 0.5mL of isopropanol for 10 minutes, the samples were again centrifuged for 10 minutes. The supernatant was removed and the RNA pellet was washed with 75% ethanol. The pellet was dissolved in RNase-free water and the quality of RNA was assessed using a NanoDrop spectrophotometer.

#### 3’ RACE assays

cDNA was generated from 1µg RNA from MiaPaCa2 as well as Suit2 cell lines using Superscript II Reverse Transcriptase (Cat# 18064022, ThermoFisher Scientific) using the primer P: 5’-GACTCGAGTCGACATCGATTTTTTTTTTTTTTTTT-3’. To PCR amplify the 3’-UTR forms of candidate genes, a gene specific forward primer spanning the stop codon of the gene was used in conjunction with a reverse primer P’: 5’- GACTCGAGTCGACATCG-3’ targeting the adapter region introduced by primer P. The PCR mixture was run on a 1.5% agarose gel and visualized using the Chemidoc imaging system followed by analysis with Image Lab software (Version 6.0.0, Bio-Rad). An identical cDNA generation and PCR procedure was followed for RNA extracted from PDA patient tumor samples. RNA from PDA patient samples were obtained from Roswell Park Pathology Shared Resource. Approval of biospecimen use was granted by the Roswell Park IRB.

#### Constructs for reporter assays

The long and short 3’-UTRs were PCR amplified from genomic DNA or BAC DNAs procured from RPCI-11 human BAC library resource at Roswell Park and subcloned into the *Renilla* luciferase vector pIS1 (Plasmid# 12179, Addgene) between the XbaI/EcoRV and NotI restriction sites. The primers were designed in accordance with 3’-UTR length estimates obtained from the 3’ RACE. The following primers were used:

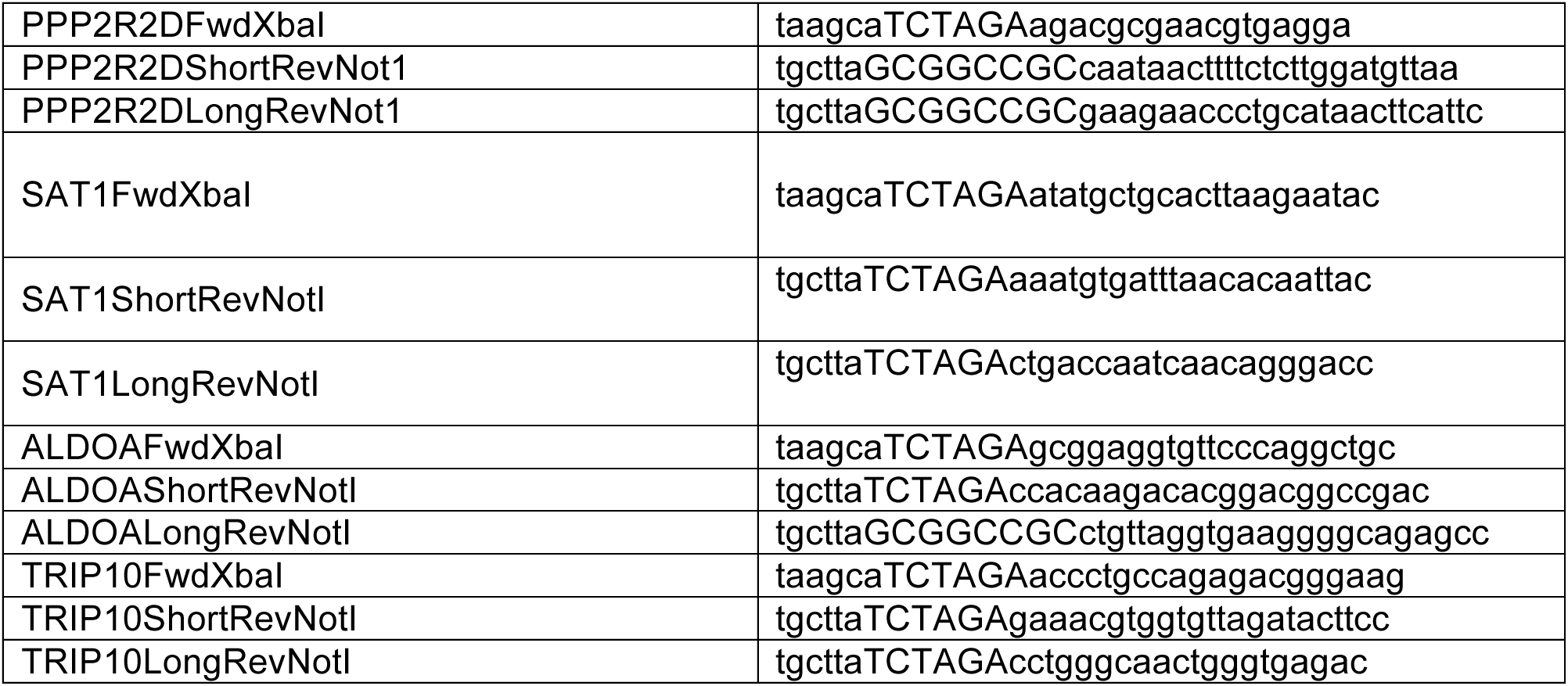
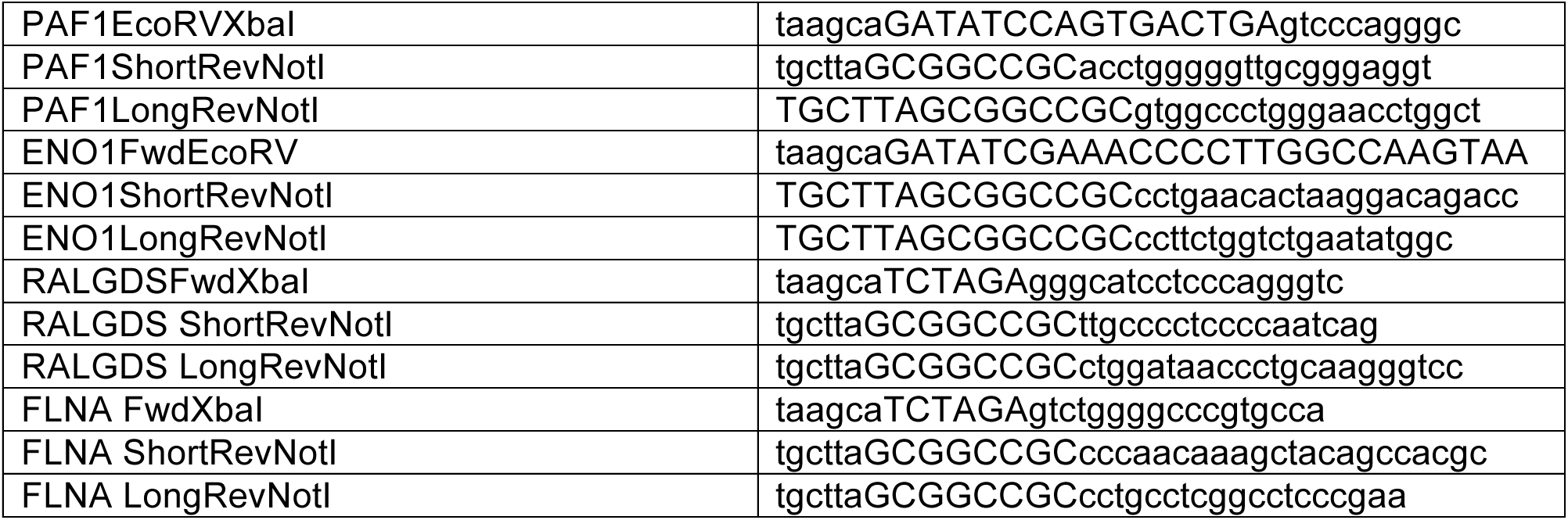

#### Luciferase reporter assays

MiaPaCa2 cells were seeded at ~10000 cells per well in a 96-well white plate (Cat# 07-200-628, Fisher Scientific). The cells were transfected the next day at ~ 60% confluency with 200ng of *Renilla* luciferase reporter plasmid (pIS1 containing the 3’-UTR region of interest) and 2ng of firefly luciferase reporter control plasmid pIS0 per well. Luciferase readings were measured 24h post-transfection with the Dual luciferase reporter assay system (Cat# E1910, Promega) using the Synergy H1 plate reader. The *Renilla* reporter reading was normalized to its corresponding firefly reading in every well to control for transfection efficiency.

#### D4476 studies

For dose-response measurements, MiaPaCa2 and Suit2 cells were seeded at a concentration of 2500 cells per well in a 96-well white plate. The next day, D4476 was titrated over a range of concentrations using the Tecan D300e Digital Dispenser and cell viability was measured 96h post drug titration using a CellTiter-Glo assay. For cell proliferation experiments, MiaPaCa2 or Suit2 cells were seeded at a concentration of 250 cells per well in a 96-well clear plate (Cat# 130188, Thermo Fisher Scientific). DMSO control or D4476 was dispensed at varying concentrations and imaged on the Cytation™ 5 Cell Imaging Multi-Mode Reader to image cell count (high contrast bright field) over time. For clonogenic experiments, MiaPaCa2 or Suit2 cells were seeded at a concentration of 250 cells per well and treated with different concentrations of D4476. The cells were allowed to grow over a period of 8-10 days after which they were fixed (10% methanol, 10% acetic acid) and stained with 0.5% crystal violet solution (in methanol). The plates were rinsed with PBS (137mM NaCl, 2.7mM KCl, 6.5mM Na2HPO4, 1.5mM KH2PO4), dried overnight and scanned. The resulting images were quantified using ImageJ (Version 1.50i). The images were uniformly thresholded and quantified for number of particles (colonies).

#### *CSNK1A1* knockdown experiments

Three different shRNAs against *CSNK1A1* gene as well as a non-targeting control shRNA (Con shRNA) were used to generate MiaPaCa2 or Suit2 control and CK1α knockdown cells. Knockdown was confirmed via immunoblotting. Briefly, samples were run alongside a molecular weight ladder (Cat# 26624, Thermo Fisher Scientific) on 10% SDS PAGE gels and then transferred to PVDF membranes (Cat# IPVH00010, Thermo Fisher Scientific) at 100V for 1 h. The membrane was blocked with 5% non-fat dry milk powder in PBST (PBS+ 0.1% Tween-20) for 1h and then incubated in the same buffer containing the primary antibody overnight on a shaker at 4°C. Polyclonal anti-CK1α (1:1000) and monoclonal β-actin (1:1000) were used to detect CK1α and β-actin respectively. The membrane was washed 4 x 5 min in PBS-T, followed by incubation with HRP-conjugated secondary antibodies (1:1000) for 1 h and then another 4 x 5 min washes. The blots were soaked with the ECL substrate (Cat# 32106, Thermo Fisher Scientific) and imaged. For cell proliferation experiments, control and CK1α knockdown Suit2 cell lines were seeded at a concentration of 250 cells per well in a 96-well white plate. Cell proliferation was measured on Day 1, 3, 5 and 7 using a CellTiter-Glo assay. The same procedure was repeated for MiaPaCa2 cells with a seed concentration of 500 cells per well. For clonogenic assays, MiaPaCa2 or Suit2 cells were seeded at a concentration of 500 cells per well in a 6-well clear plate. The cells were allowed to grow over a period of 8-10 days, fixed, stained and quantified as described previously.

#### Statistical analyses

All findings presented were replicated in three or more independent experiments. Comparisons between two groups were performed using unpaired *t-test* with Welch’s correction in Graph Pad Prism 8. In general, p < 0.05 was considered significant, and the determined p values are provided in the figure legends. Asterisks in graphs denote statistically significant differences as described in figure legends.

