## Supplementary Figures for "Alternative polyadenylation drives oncogenic gene expression in pancreatic ductal adenocarcinoma"

Venkat S, *et al.*, Figure S1

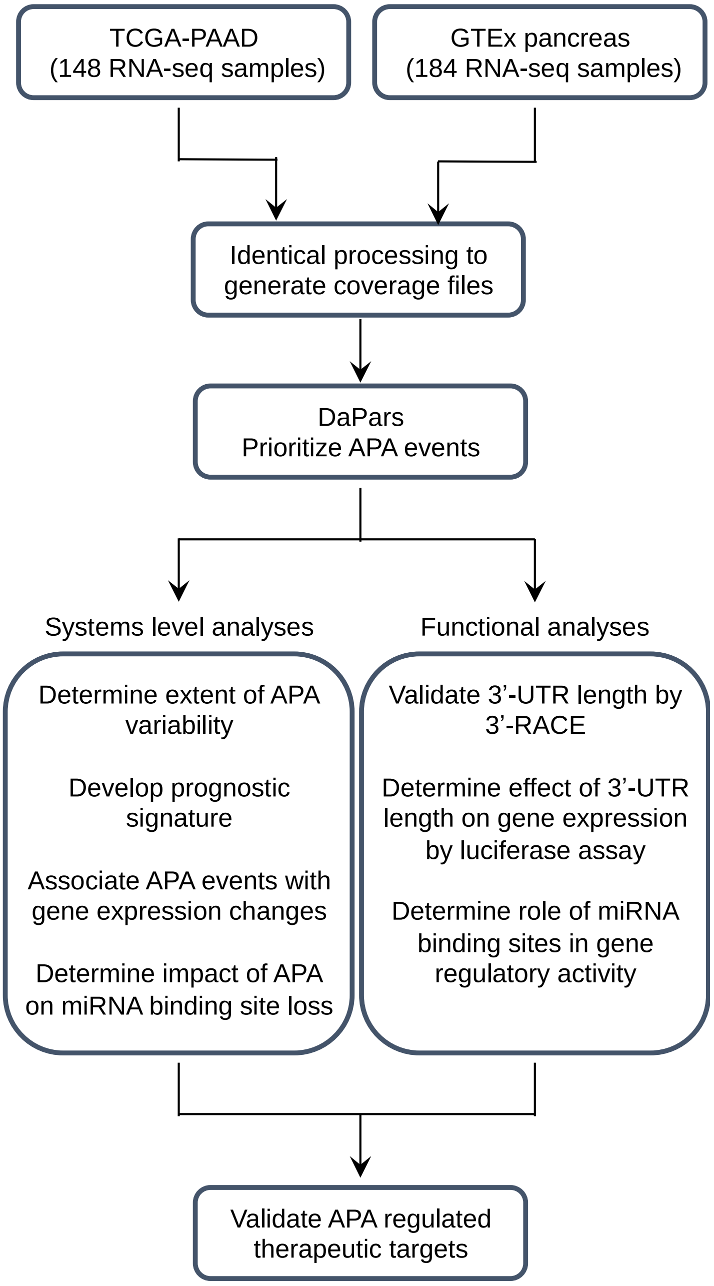

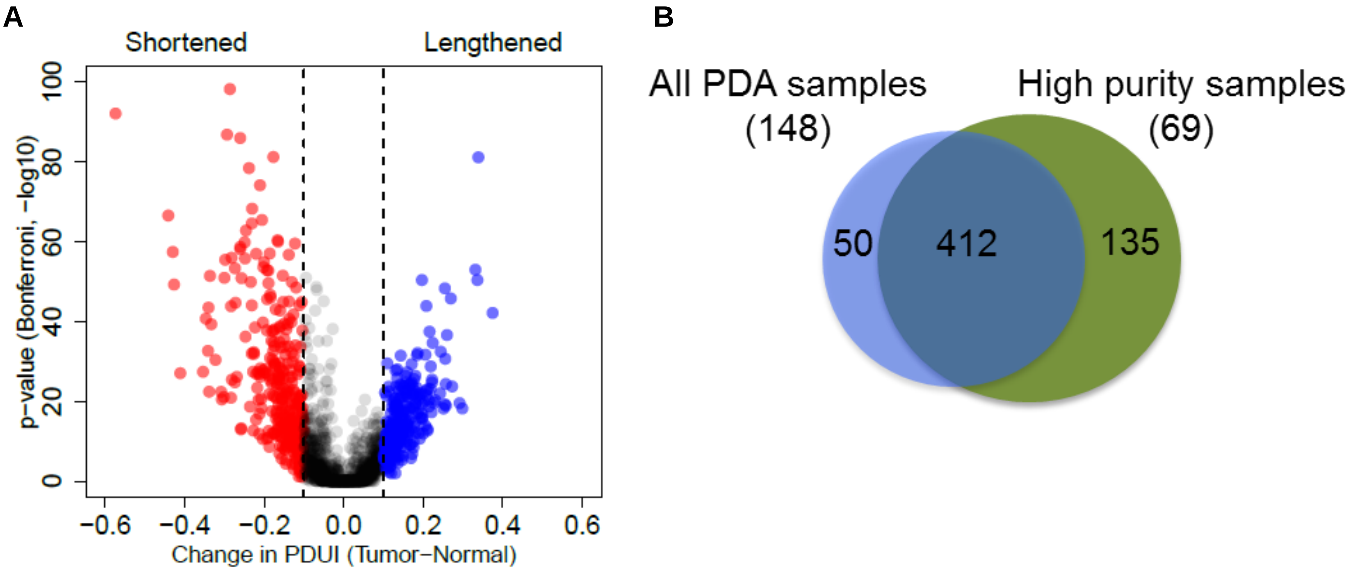


**C**

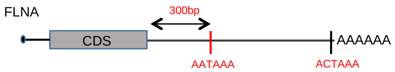


**TGA**gtctggggcccgtgccagccggcagcccccaagcctgccccgctacccaagcagccccgccctcttcccctcaaccccggcccaggccgccctggccgcccgcctgtcactgcagccgcccctgccctgtgccgtgctgcgctcacctgcctccccagccagccgctgacctctc
ggctttcacttgggcagagggagccatttggtggcgctgcttgtcttctttggttctgggaggggtgagggatgggggtcctgtacacaaccacccactagttctcttctccagccaagagg **aataaa**gttttgcttccattctcctttgcgtggctgtagctttgttggggttgggggaccagtcaagg
ccaacctgacaggggtgccaatgagcagatcaaagcatgggctgagcacaggtcacctccctccagggctcctgcttcaccccaccctgactccagaaggcaccatctatctttgcttcctctgcctccctgagttcaagccaacgtaaggagaggaacatgtagcaacctggaatggag
cccaggcagcaaggaggcaaagagctgggcaggagagcccattactgggtagggtggtgtcaggaaggctccctggtggaggtggccagataatcaacagcaggcccggccgggtgtggtggctcacgcctgtaatcccagcacttcgggaggccgaggcaggcagatcacaag
gtcaggagatcaagaccatcctggctaacacggtgaaaccccgtctct **actaaa**aatacaaaaaattagccaggtgtggtggcgggcgcctgtagtcccagctactcgggaggctgaggcaggagaatggtgcgaacccaggagacagagcttgcagtgagccgagatcgcgccac
tgccctccagcctgggcgacagagcgagactctgtctcaaaaaaaaaaaaaaaaaaaaaatcacagcaggccctggccatctgccaggtaacttgaacatgccaccatgccaccattcccccaaacaagggactgggatgaggccccctcccccaggctctgggcctcagggcccttt
agttaacaggtggcaggaggtggatgcattcaaggtgccaatcccaggtaccctgggccccctgctgaatatcccctcattggtaccctgccgtggattcccacaccaacctatgtgtctcccactgtaaagattgcccccaaacacataacatcccaagtcgtcctggctcctggtgctgcct
gcatgctatcgggacatcccaactgcagcacccaaagctgcctgctccctgggccaaagg

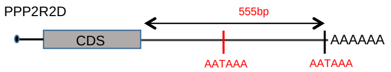


**TAG**agacgcgaacgtgaggaccaagtcttgtcttgcatagttaagccggacatttttctgtcagagaaaaggcatcattgtccgctccattaagaacagtgacgcacctgctacttcccttcacagacacaggagaaagccgcctccgctggaggcccggtgtggttccgcctcggcga
ggcgcgagacaggcgctgctgctcacgtggagacgctctcgaagcagagttgacggacactgctcccaaaaggtcattactcag **aataaa**tgtatttatttcagtccgagccttcctttccaatttatagaccaaaaaattaacatccaagagaaaagttattgtcagataccgctctttctcc
aactttccctctttctctgccatcacacttgggccttcactgcagcgtggtgtggccaccgtccgtgtcctctcggccttcctccgagtccaggtggactctgtggatgtgtggatgtggcccgagcaggctcaggcggccccactcacccacagcatccgccgccacccccttcgggtgtgagc
gctc**aataaa**aacaacacactataaagtgtttttaaatccaaacagaagtattgtctttttatttaattttattgcatagaatgaagttatgcagggttcttctttggaactaactgtttgagaaatgtgtgtcct

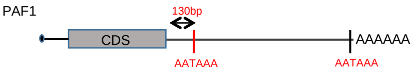


**TGA**gtcccagggcattcagggctggttcagacaccattattgtgagcagcaaagcacttttctagtggtctgtttgtgagcctttcacttgtttgttccccacccccaaacctttgctgtt **aataaa**gtcaacttctctttacttcccctcccagcccctatgatgtcattcagcccagcacatacctcc
cgcaacccccaggtgctcaaggatcttggcttgaggcttagaaacacaggccagaggcagggccaatggtttcccacaaaatgtagcacaacgaaggtcagaattcgtttttttgtttgtttttttttttccaaaataagcccagaccattaaacaagtgaaactccaacaaataagtcttctc
caacagcgagaaaaactgtacagttactcaaagctgattctgccagtggggccggggacagaagtgggtagggagggtgaaatcatggaggagggcctggggagggggctggagcgggagagggtcagggtcctgcccatcagagtggggccgcctgcgtcctgcacactctg
ctgtcaggtggggtgggggcagctcttgcctccctgtgtgtgtgagacggtgtccctcaccacctcccagtgccaggcacagtcagctcaccctgatggagagtgaacaagacaggacaggctggggagaggggagggtcccatgagagggacaccctgctgattccaaacgagg
tggcccatccatcccacttcctgcagctagtgggagggggactggagaagccctgggagattgaggggatgagcaaaggcacagaaacatggagggccggggtgggggacttgcataggttgatgagtgtgcaagaggtacata **aataaa**cctgacaccccacccagcccggc
actgggagaggaagtggggacccagagccaggttcccagggccac

**Venkat S, *et al.*, Figure S3**

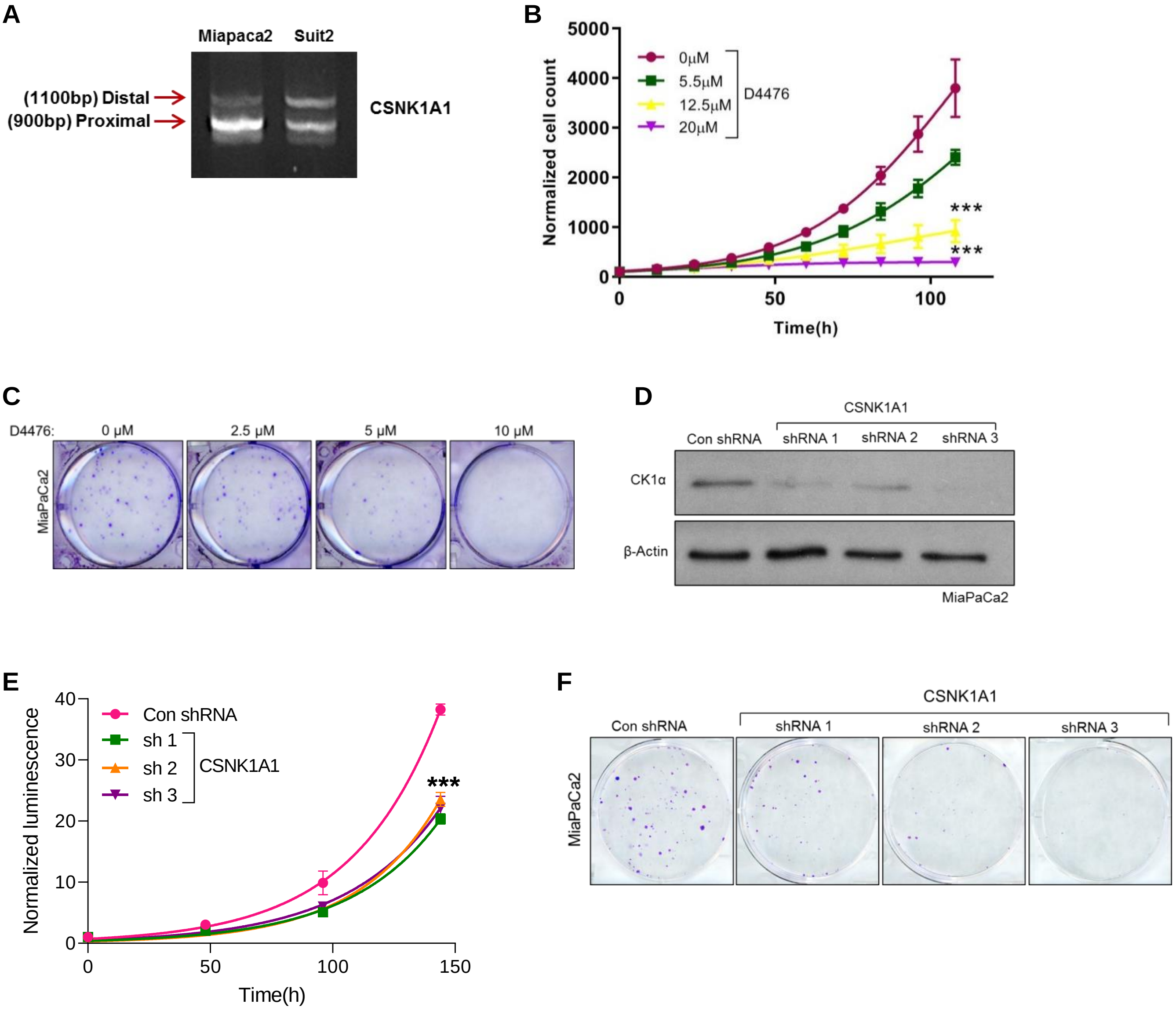

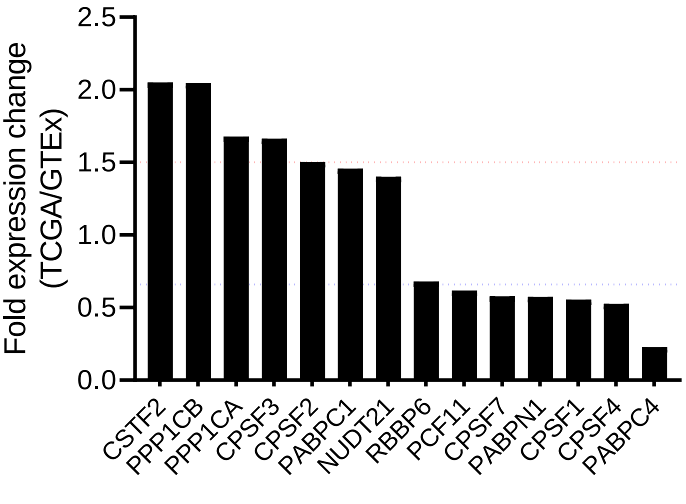
Fold expression change
(TCGA/GTEx)
2.5
2.0
1.5
1.0
0.5
0.0
CSTF2
PPP1CB
PPP1CA
CPSF3
CPSF2
PABPC1
NUDT21
RBBP6
PCF11
CPSF7
PABPN1
CPSF1
CPSF4
PABPC4
